# Functional microstructure of Ca_V_-mediated calcium signaling in the axon initial segment

**DOI:** 10.1101/2020.11.13.382374

**Authors:** Anna M Lipkin, Margaret M Cunniff, Perry WE Spratt, Stefan M Lemke, Kevin J Bender

## Abstract

The axon initial segment (AIS) is a specialized neuronal compartment in which synaptic input is converted into action potential output. This process is supported by a diverse complement of sodium, potassium, and calcium channels (Ca_V_). Different classes of sodium and potassium channels are scaffolded at specific sites within the AIS, conferring unique functions, but how calcium channels are functionally distributed within the AIS is unclear. Here, we utilize conventional 2-photon laser scanning and diffraction-limited, high-speed spot 2-photon imaging to resolve action potential-evoked calcium dynamics in the AIS with high spatiotemporal resolution. In mouse layer 5 prefrontal pyramidal neurons, calcium influx was mediated by a mix of Ca_V_2 and Ca_V_3 channels that differentially localized to discrete regions. Ca_V_3 functionally localized to produce nanodomain hotspots of calcium influx that coupled to ryanodine-dependent stores, whereas Ca_V_2 localized to non-hotspot regions. Thus, different pools of Ca_V_s appear to play distinct roles in AIS function.

## Introduction

Voltage-gated calcium channels (Ca_V_s) occupy a unique functional niche in neurons, affecting both electrical signaling across the membrane and initiating intracellular cascades that regulate ion channel function, cellular processes, and gene expression. Ca_V_s are distributed broadly across somatodendritic and axonal compartments, but only recently have we come to appreciate their role at the intersection of these two compartments, the axon initial segment (AIS). The AIS is enriched with sodium and potassium channels scaffolded by a complex intracellular skeleton and can be a site for specialized inhibitory synaptic input (Bender and Trussell, 2012; Huang and Rasband, 2018; Kole and Stuart, 2012; Leterrier, 2018). Of all Ca_V_ classes, low voltage-activated Ca_V_3s appear to be most commonly expressed in the AIS. AIS Ca_V_3 channels were first shown to regulate the threshold and timing of APs in auditory brainstem cartwheel interneurons, somatosensory cortex pyramidal cells, and cerebellar Purkinje cells (Bender and Trussell, 2009; Bender et al., 2012). AIS-localized Ca_V_3 channels have also been identified at the AIS of cells in avian brainstem and murine cerebellum, hippocampus, and frontal cortex (Clarkson et al., 2017; Dumenieu et al., 2018; Fukaya et al., 2018; Hu and Bean, 2018; Jing et al., 2018; Martinello et al., 2015). In many of these cells, Ca_V_3 channels appear to be interspersed in the AIS with other Ca_V_ classes. This diversity is most apparent in neocortical pyramidal cells, where calcium influx has been reported to be mediated by a range of channel types, including Ca_V_1, members of the Ca_V_2 family, and Ca_V_3 (Clarkson et al., 2017; Hanemaaijer et al., 2020; Yu et al., 2010)

Across neuronal compartments, the spatial organization of Ca_V_s shapes function by linking spatially-restricted calcium influx to nearby calcium-sensitive processes. In somatodendritic compartments, coupling of Ca_V_s to calcium-activated potassium channels regulates action potential (AP) dynamics (Bock and Stuart, 2016; Irie and Trussell, 2017), EPSP amplitude and summation (Chen-Engerer et al., 2019; Wang et al., 2014), and calcium influx (Jones and Stuart, 2013). In the soma, calcium influx through Ca_V_1 channels influences activity-dependent transcription by nuclear transcription factors through interactions with calmodulin factors (Wheeler et al., 2012). And in axon terminals, the density and spatial arrangement of Ca_V_s relative to neurotransmitter release machinery determines release probability (Rebola et al., 2019; Scimemi and Diamond, 2012), shaping both the dynamics of short-term plasticity and its regulation by neuromodulators (Bucurenciu et al., 2008; Burke et al., 2018; Vyleta and Jonas, 2014). However, the organization of Ca_V_s d within the AIS, and how they interact with calcium sensitive signaling pathways, remains unclear.

The AIS serves multiple roles, acting both as a site of electrogenesis for APs as well as a diffusion barrier between somatodendritic and axonal compartments (Bender and Trussell, 2012; Leterrier and Dargent, 2014). These functions are supported by a complex scaffold of intracellular and membrane-bound proteins. Rings of actin connected by spectrins occur periodically along the AIS, forming a scaffold for ankyrin-G to bind and anchor voltage-gated sodium channels (Na_V_) and voltage-gated potassium (K_V_) K_V_7 channels (Leterrier, 2018). K_V_1.1 and K_V_1.2 are anchored by a complex that includes PSD-93 the paranodal protein Caspr and typically cluster at actin rings (Ogawa and Rasband, 2008; Pinatel et al., 2017). K_V_2.1 channels are found at yet another clustering domain enriched with the scaffolding protein gephyrin (King et al., 2014). In neocortical pyramidal cells, these gephyrin-rich sites are punctate, occupying small gaps in an otherwise continuous sheath of Na_V_-rich membrane. It is here that chandelier cells form GABAergic synapses at the AIS (Inan and Anderson, 2014). Furthermore, a specialized endoplasmic reticulum, termed the cisternal organelle, abuts these gephyrin-rich regions. These cisternal organelles express ryanodine receptors (RyRs) (King et al., 2014), which mediate calcium-induced calcium release from intracellular stores. Interestingly, we have previously shown that RyR-dependent signaling is necessary for dopaminergic signaling cascades that regulate AIS Ca_V_3 function (Yang et al., 2016), but the local calcium source that evokes calcium-induced-calcium-release at these sites has not been identified. Given the differential distribution of other ion channel classes in the AIS, we hypothesized that a unique complement of Ca_V_s may be localized to these regions and engage RyRs.

Here, we developed diffraction-limited, high-frequency 2-photon imaging techniques to explore the functional microstructure of AP-evoked calcium signaling in mouse prefrontal pyramidal cell initial segments. We found that calcium influx was mediated by a mix of Ca_V_2.1, 2.2, 2.3 and Ca_V_3-type calcium channels that were distributed into distinct functional domains. In some regions, micron-wide “hotspots” of fast, high-amplitude calcium influx occurred. These hotspots were dominated by Ca_V_3-mediated calcium influx, whereas non-hotspot regions were dominated by Ca_V_2.1/2.2-mediated influx. Furthermore, Ca_V_3 channels were preferentially linked to RyR-dependent intracellular stores, suggesting that AIS Ca_V_3 channels, commonly expressed in many neuronal classes, are complexed in pyramidal cells to regions associated with GABAergic synaptic transmission. Thus, Ca_V_s may play distinct roles in different subcompartments within the AIS.

## METHODS

### Ex vivo electrophysiological recordings

All experiments were performed in accordance with guidelines set by the University of California Animal Care and Use Committee. C57Bl/6 mice of both sexes aged P20 through P30 were anesthetized and 250 μm coronal sections containing medial prefrontal cortex were collected. Cutting solution contained, in mM: 87 NaCl, 25 NaHCO_3_, 25 glucose, 75 sucrose, 2.5 KCl, 1.25 NaH_2_PO_4_, 0.5 CaCl_2_, and 7 MgCl_2_, bubbled with 5% CO_2_/95% O_2_. After cutting, slices were incubated in the same solution for 30 min at 33°C, then at room temperature until recording. Recording solution contained, in mM: 125 NaCl, 2.5 KCl, 2 CaCl_2_, 1 MgCl_2_, 25 NaHCO_3_, 1.25 NaH_2_PO_4_, and 25 glucose, bubbled with 5% CO_2_/95% O_2_. Recordings were done at 32-34°C, with the exception of SBFI and Fluo4ff experiments, which were performed at room temperature (22°C). Osmolarity of the recording solution was adjusted to ∼310 mOsm.

Neurons were visualized using Dodt contrast optics for visually-guided whole-cell recording. Patch electrodes were pulled from Schott 8250 glass (3-4 MΩ tip resistance) and filled with a solution containing, in mM: 113 K-gluconate, 9 HEPES, 4.5 MgCl_2_, 14 Tris_2_-phoshocreatine, 4 Na_2_-ATP, 0.3 Tris-GTP, ∼290 mOsm, pH 7.2-7.25. Calcium buffers, volume filling dyes, and calcium or sodium indicators were included in the internal solution as follows: for linescan calcium imaging experiments, 250 μM Fluo-5F and 20 μM Alexa 594 were added. For fast (5.3kHz) linescan sodium imaging, 2 mM SBFI, 0.1 μM EGTA, and 20 μM Alexa 594 were added. For fast linescan calcium imaging, 500 μM Fluo-4FF and 0.1 μM EGTA were added. For calcium imaging at single diffraction limited spots, 600 μM OGB-5N, 0.1 μM EGTA and 20 μM Alexa 594 were added. For pointscan sodium imaging, 500 μM ING-2 was added.

Electrophysiological data were acquired using a Multiclamp 700B amplifier (Molecular Devices). For fast linescan experiments, data were acquired at 50 kHz and filtered at 20 kHz. For all other experiments, data were acquired at 20 kHz and filtered at 10 kHz. All recordings were made using a quartz electrode holder to eliminate electrode drift within the slice, enabling stable imaging of diffraction-limited spots in close proximity to the recording electrode (Sutter Instruments). Cells were excluded if series resistance exceeded 20 MΩ or if the series resistance changed by greater than 30%. All recordings were made from Layer 5b pyramidal neurons in prefrontal or primary somatosensory cortex and data were corrected for a 12 mV junction potential.

### Two Photon Imaging

Two photon laser scanning microscopy (2PLSM) was performed as described previously (Bender and Trussell, 2009). A Coherent Ultra II laser was tuned to 810 nm for morphology and calcium imaging and ING-2 based sodium imaging. The laser was tuned to 790 nm for SBFI-based imaging. Fluorescence was collected with either a 40x, 0.8 NA objective (data in Figs. 1-3) or a 60x, 1.0 NA objective (data in Figs. 4-6) paired with a 1.4 NA oil immersion condenser (Olympus). Dichroic mirrors and band-pass filters (575 DCXR, ET525/70 m-2p, ET620/60 m-2p, Chroma) were used to split fluorescence into red and green channels unless otherwise specified. HA10770-40 photomultiplier tubes (PMTs, Hamamatsu) selected for >50% quantum efficiency and low dark counts captured green fluorescence (Fluo-5F, Fluo-4FF). Red fluorescence (AlexaFluor 594) was captured using R9110 PMTs. For ING-2 based imaging, the epifluorescence filters were removed and the transfluorescence filters were replaced with a single 535/150 bandpass filter (Semrock) and all fluorescence was collected on HA10770-40 PMTs.

**Figure 1.**
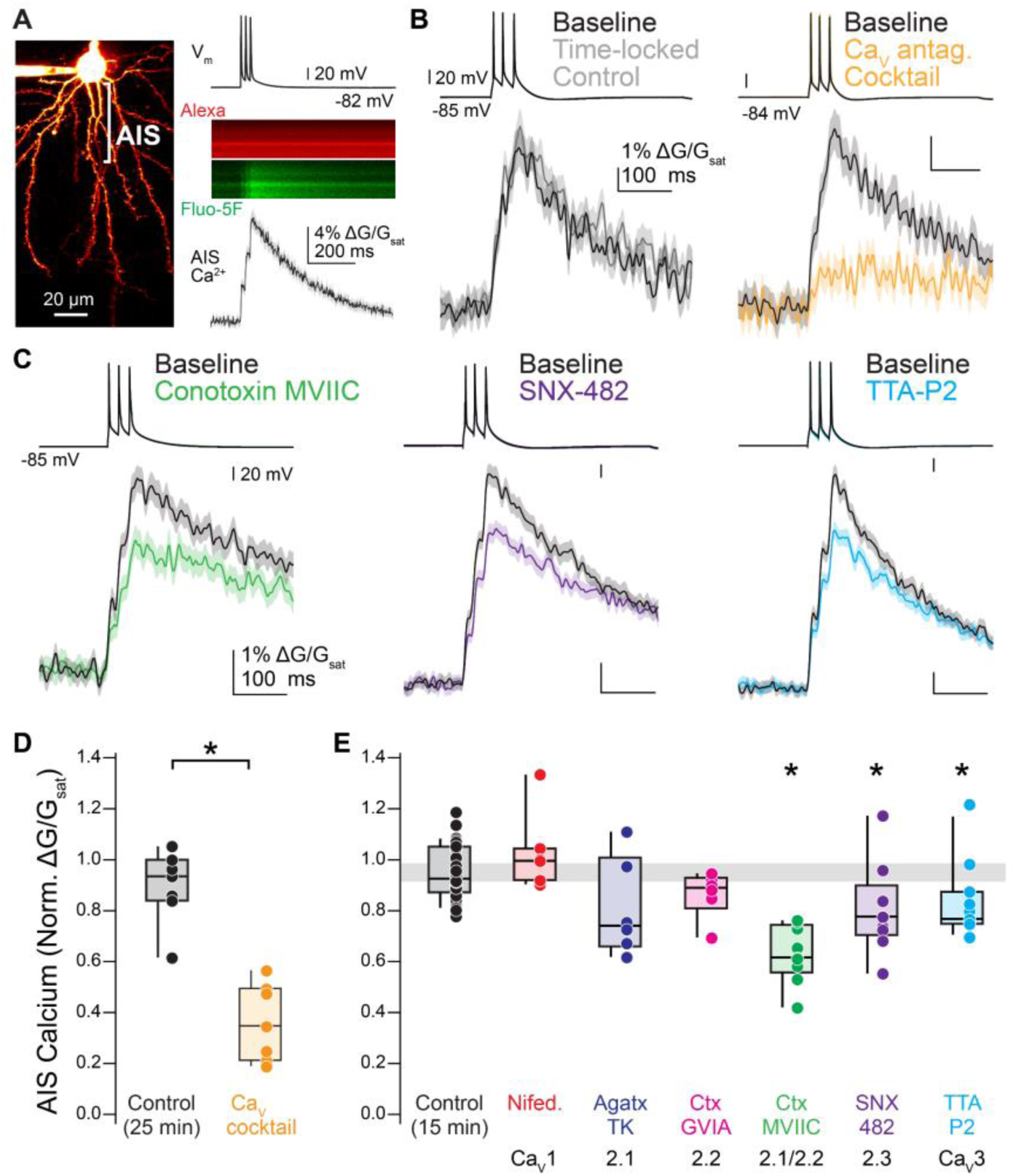
Ca_V_2.1, Ca_V_2.2, and Ca_V_3 contribute to calcium influx at the axon initial segment. A. Left: Two photon laser-scanning microscopy (2PLSM) z-stack of a Layer 5 pyramidal neuron visualized with Alexa 594. AIS indicated by bracket. Right: example linescan of AIS averaged over 20 trials. APs were evoked with somatic current injection (1 nA, 5 ms, 20 ms interstimulus interval). B. Left: Representative time-locked control cell. Linescan data displayed as mean ± standard error. Baseline, black; post, gray. Right: Representative effects of Ca_V_ antagonist cocktail on AIS calcium. Baseline, black; cocktail, yellow. C. Representative examples of selective Ca_V_ antagonists on AIS calcium. Baseline, black; antagonists, other colors. D. Summary of the effects of the Ca_V_ antagonist cocktail on AIS calcium. E. Summary of the effects of individual Ca_V_ antagonists on AIS calcium. Gray bar represents 95% confidence interval of control data.

**Figure 2.**
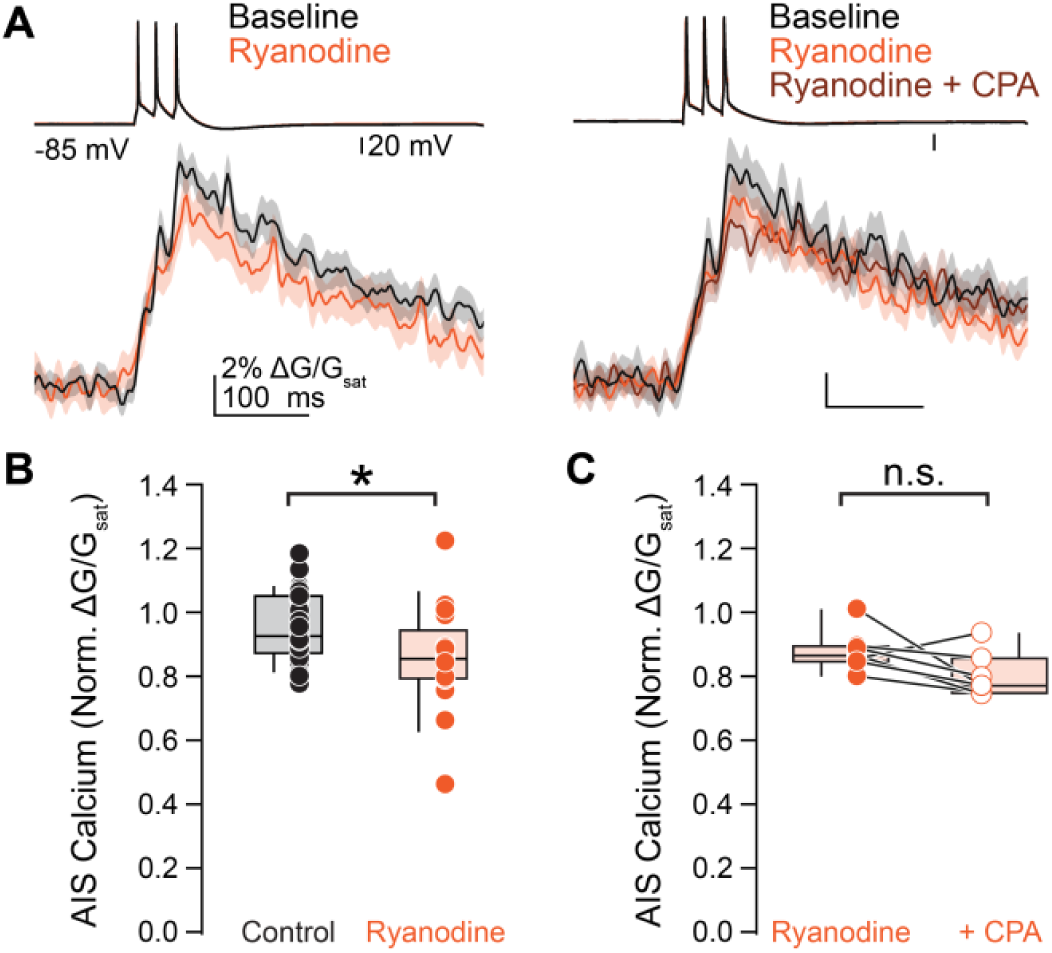
Calcium stores contribute to AIS calcium during AP firing. A. Left: Representative effect of ryanodine (20 µM) on AIS calcium. Right: Representative effect of sequential ryanodine and cyclopiazonic acid (CPA, 20 µM) application on AIS calcium. Linescan data presented as mean ± standard error. B. Summary of the effects of ryanodine on AIS calcium. C. Summary of the effects of sequential application of ryanodine and cyclopiazonic acid.

**Figure 3.**
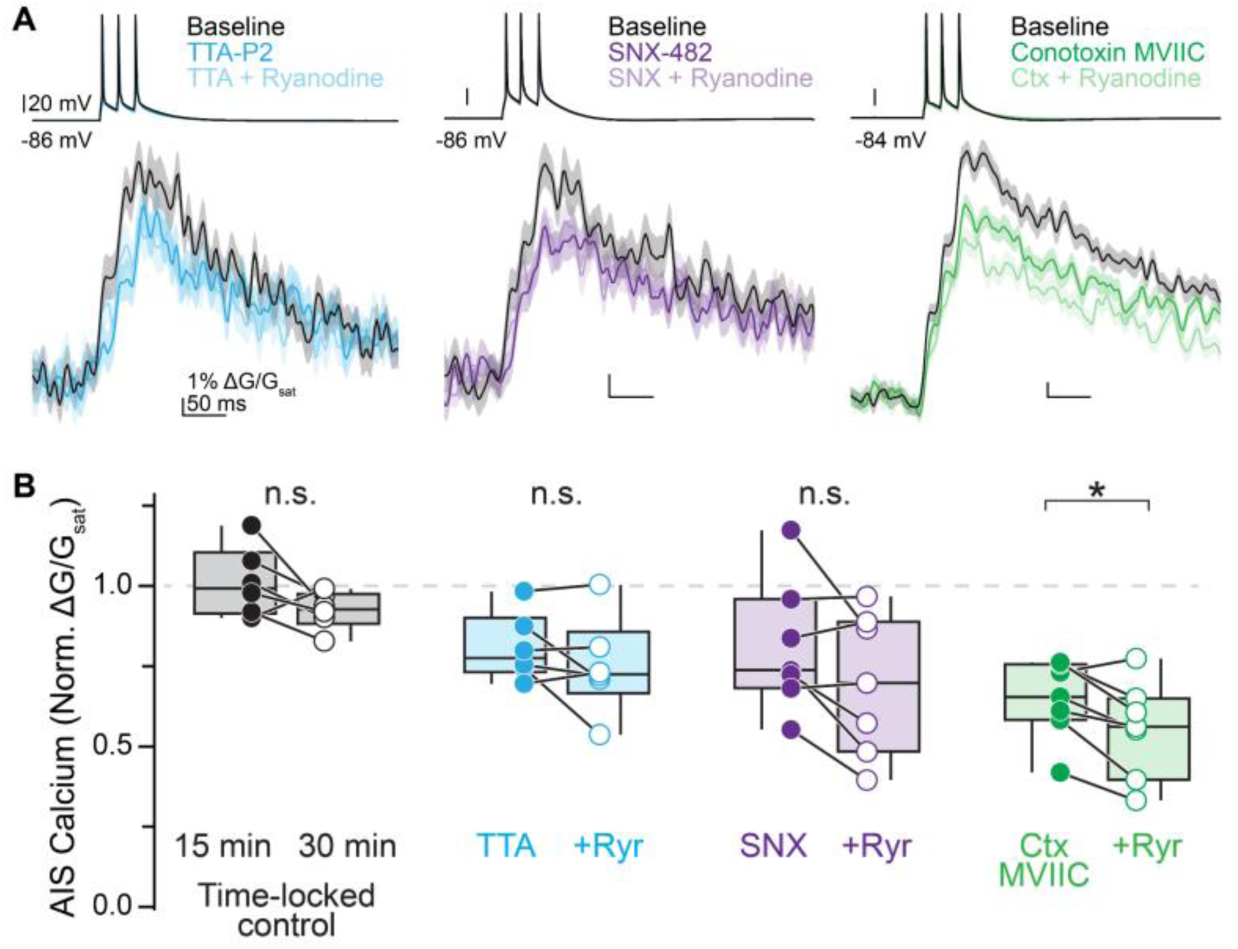
Ca_V_3 channels couple to ryanodine receptors on calcium stores. A. Representative effects of sequential block of individual Ca_V_ types and release from calcium stores. Linescan data shown as mean ± standard error. B. Summary of the effects of Ca_V_ antagonists and ryanodine block. n.s.: not significant.

**Figure 4.**
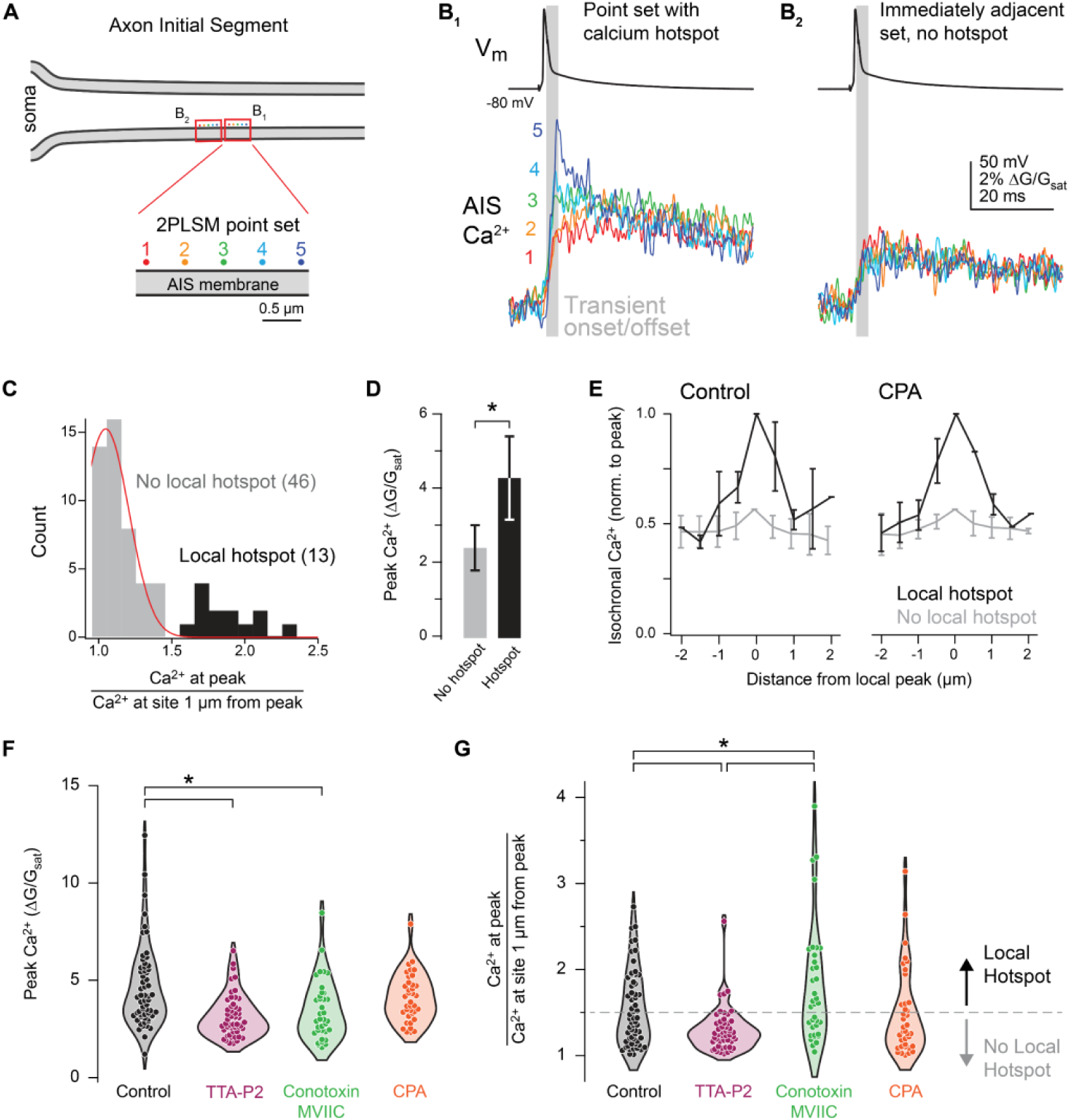
Ca_V_3 channels and Ca_V_2.1/2.2 exhibit distinct functional distributions. A. Schematic of 2PLSM point scan imaging protocol. Points were imaged in sets of 5, with each point separated by 0.5 µm. The laser was parked at a single diffraction-limited point for 25 ms preceding and 100 ms following an AP and calcium influx was measured with OGB-5N. Points were scanned in the sequence 2, 4, 1, 3, 5 and each point sampled a single AP. Data was averaged over 20-50 repetitions. B. Isochronal calcium peaks from neighboring point sets. Calcium influx at each point is color-coded as in panel A. B1 shows a point set with a hotspot at point 5. B2 is the point set immediately adjacent to B1 and shows equivalent calcium influx across all points. Gray bar indicates the calcium transient onset and offset. C. Distribution of point sets containing hotspots. Peak calcium influx at the brightest point was divided by the isochronal calcium influx at the point(s) 1 µm away. 46 of 59 sites imaged fell within a normal distribution, while 13 sites exhibited higher relative calcium influx. Black, point sets containing a local hotspot; gray, point sets with no local hotspot. Red line indicates the fit of a normal distribution. Total distribution fit for normality (Shapiro Wilk test p = 0.0016). D. Calcium influx at hotspots was approximately 2x higher than calcium influx at non-hotspot points. Black, point sets containing a local hotspot; gray, point sets with no local hotspot. Data are plotted as mean ± standard deviation. E. Comparison of the flanks of point sets with a local hotspot and those without. Black, point sets containing a local hotspot; gray, point sets with no local hotspot. Data are plotted as mean ± standard deviation for each 0.5 µm increment from the brightest point of the set. F.Influence of selective Ca_V_ antagonists or store depletion on peak calcium influx during point scan imaging. Circles represent single point sets. Black, control; pink, TTA-P2; green, ω-conotoxin MVIIC; orange, CPA. G. Influence of selective Ca_V_ antagonists or store depletion on calcium hotspots. Hotspots were classified as points >1.5 times brighter than the point(s) 1 µm away. Dotted gray line represents the distinction between point sets with a local hotspot (above) and those without (below). Color code as in panel F.

**Figure 5.**
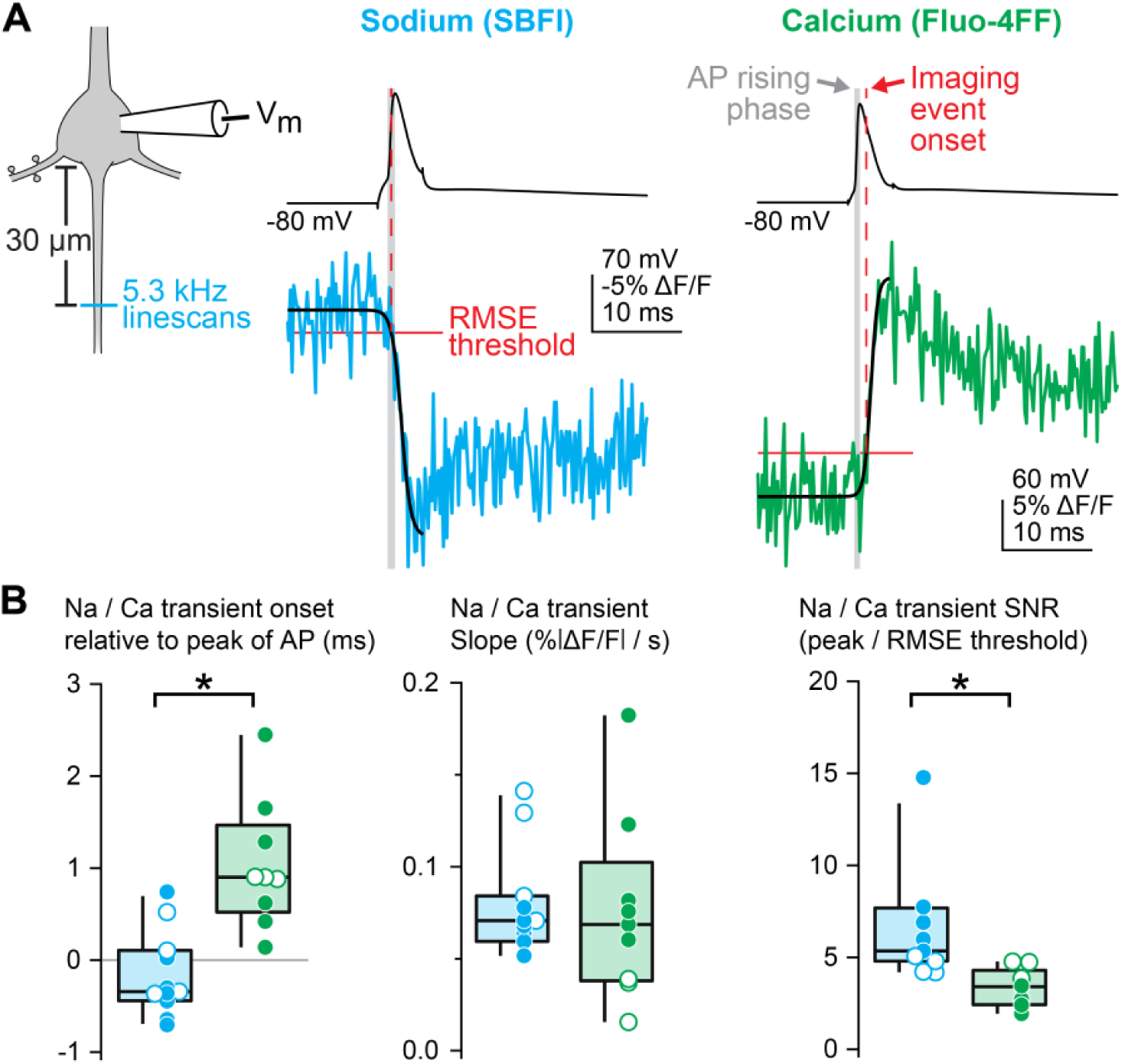
Fast linescan imaging reveals distinct temporal profiles of sodium and calcium influx. A. Schematic of fast linescan protocol. Left: Linescan imaging was performed transecting the AIS at 5.3 kHz with either the sodium dye SBFI or the calcium dye Fluo-4FF. Middle: representative example of sodium influx aligned to soma-evoked AP shows that sodium influx begins during the rising phase of the AP. Right: representative example of calcium imaging aligned to AP. Calcium influx occurs during AP repolarization. Blue, SBFI; green, Fluo-4FF. Dashed vertical red line indicates imaging event onset. Gray bar represents rising phase of the AP (threshold to peak). Black line is the sigmoid fit of the linescan. Solid red line shows baseline noise of the linescan. B. Comparison of sodium and calcium transient onset time, slope, and signal-to-noise ratio. Left: summary of the timing of sodium and calcium transients relative to the peak of the AP. Negative values represent transient onset that precedes the AP peak. Middle: The slope of optically-recorded sodium and calcium transients. Slope was calculated using the sigmoid fit. Right: signal-to-noise ratio for sodium and calcium transients. Circles represent individual cells. Open circles are cells with similar signal-to-noise ratios. Blue, SBFI; green, Fluo-4FF.

**Figure 6.**
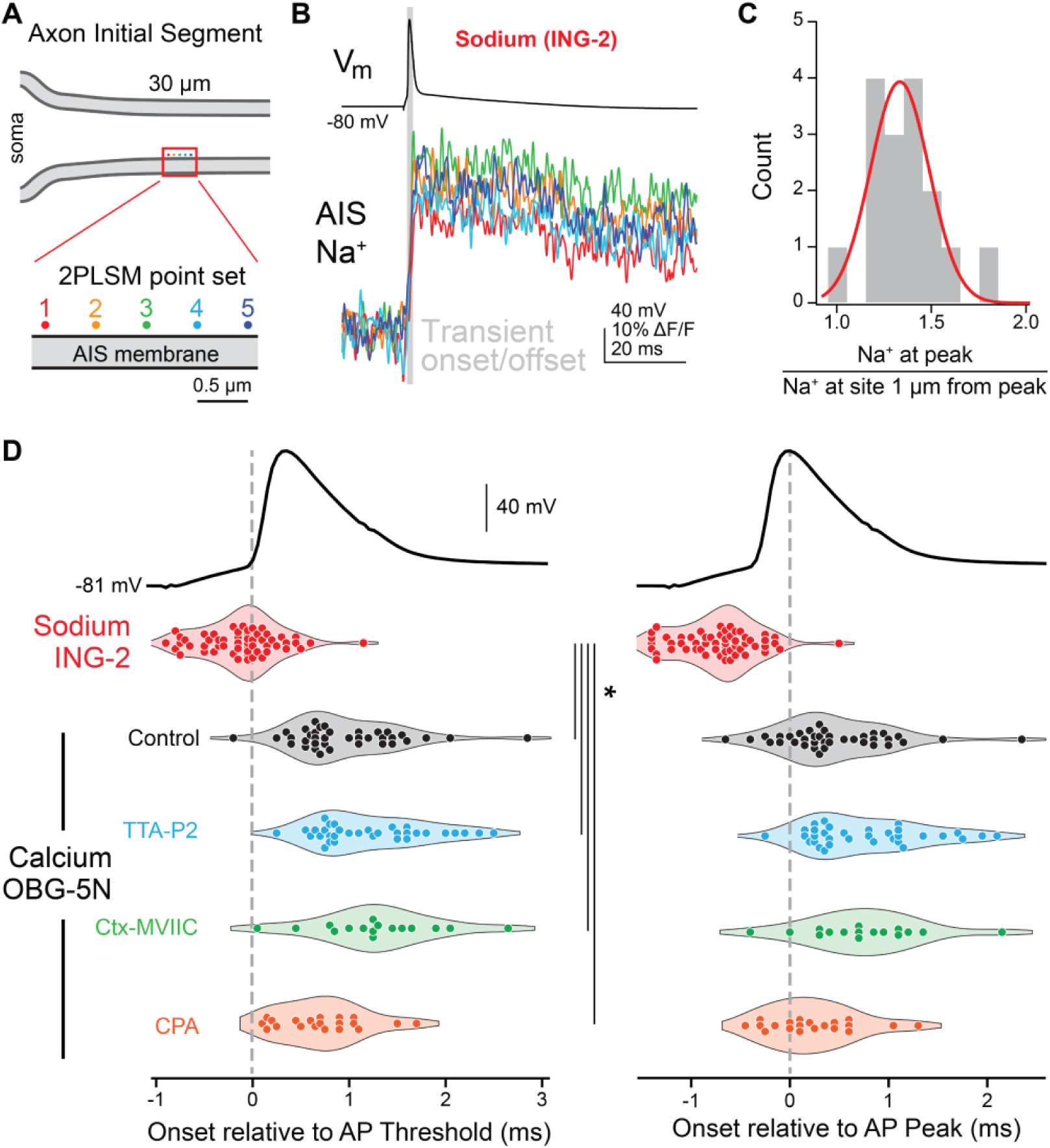
Sodium and calcium influx occur on the rising and falling phases of the AP at physiological temperatures, respectively. A. Pointscan imaging protocol was performed as in Figure 4A. OGB-5N was replaced with the sodium indicator ING-2, and Alexa-594 was excluded from the internal recording solution. B. Representative ING-2 sodium imaging pointset. Points are color-coded as in Panel A. Gray bar indicates the sodium transient onset and offset. C. Distribution of sodium imaging point sets calculated as in Fig 4C. Red line represents the fit of a normal distribution to the data. D. Sodium and calcium transients from pointscan imaging temporally-aligned to AP threshold and peak. Left: Sodium and calcium influx onset relative to AP threshold. Right: Sodium and calcium influx onset relative to AP peak. Transient onset time was measured for the brightest point in a point set. Circles are individual point sets. Gray dashed line shows AP threshold (left) or peak (right) timing. Red, ING-2 sodium imaging; black, OGB-5N calcium imaging in control conditions; blue, calcium imaging in the presence of TTA-P2; green, calcium imaging in the presence of ω-conotoxin MVIIC; orange, calcium imaging in the presence of cyclopiazonic acid.

Fluorescence data were collected either using linescan or pointscan configurations. In linescan mode, the laser was repeatedly scanned over a region of axon at a rate of ∼0.5 or 5.3 kHz. For 0.5 kHz calcium imaging, data were averaged over 20-40 trials and reported as ΔG/G_sat_, which was calculated as Δ(G/R)/(G/R)_max_*100 where G/R_max_ is the maximal fluorescence in saturating calcium (2 mM). For 5.3 kHz imaging, data were averaged over 50 trials and reported as the change in fluorescence detected by HA10770-40 PMTs (ΔG/G). In pointscan mode, the laser was parked at a single diffraction-limited spot and calcium and sodium influx were imaged with OGB-5N and ING-2, respectively, for 25 ms preceding and 100 ms following an AP. Fluorescence data were acquired at 20 kHz. Points were imaged in sets of 5, each sampling a single AP, spaced at 0.5 µm intervals along the visualized edge of the axon. Individual points were imaged in a sequence of 2,4,1,3,5, with 2 being the point most proximal to the soma. Individual APs within the set of 5 points were separated by 250 or 500 ms for calcium and sodium imaging, respectively. Data were averaged over 20-50 repetitions and then smoothed using a 40 point binomial filter in IgorPro before analysis.

### Chemicals

TTA-P2 was from Alomone Labs. ω-conotoxin-MVIIC, ω-conotoxin-GVIA, ω-agatoxin-TK, and SNX-482 were from Peptides International. Nifedipine was from Tocris. All calcium channel antagonists were prepared as stock solutions in ddH20 in glass vials. Ryanodine was from Tocris and was prepared as a stock solution (25 mM) in DMSO (0.08% final concentration DMSO). Peptide toxins were applied in recording solution supplemented with 1% bovine serum albumin to minimize peptide pre-absorption. Recording solution reservoirs and tubing connecting the reservoir to the recording chamber were made of borosilicate glass, except for 30 mm lengths of Tygon tubing fed through the recirculation peristaltic pump (Ismatec Reglo). Alexa Fluor 594 hydrazide Na^+^ salt, Fluo-5F pentapotassium salt, SBFI tetraammonium salt, Fluo-4FF pentapotassium salt, and Oregon Green 488 BAPTA-5N hexapotassium salt were from Invitrogen. ION NaTRIUM-Green-2 TMA+ salt (ING-2) was from Abcam.

### Statistics

All data are reported as medians with inter-quartile ranges in text and displayed with box plots (medians, quartiles and 90% tails) or violin plots with individual data points overlaid. For linescan experiments, n denotes cells. For pointscan experiments, n denotes point sets, and the number of cells is reported in the text. For cells in Fig. 1-3, time-locked control cells were interleaved with antagonist cells. Sample sizes were chosen based on standards in the field. No assumptions were made for data distributions, and unless otherwise noted, two-sided, rank-based nonparametric tests were used. Significance level was set for an alpha value of 0.05, and a Holm-Sidak correction was used for multiple comparisons when appropriate. Statistical analysis was performed using Statview, IgorPro 8.0, and the Real Statistic Resource Pack plugin for Microsoft Excel (Release 7.2).

## RESULTS

While action potential-evoked Ca_V_-mediated calcium influx has been observed in the AIS of a range of cell classes, the channels that mediate such influx appear to vary from class to class (Bender and Trussell, 2009; Clarkson et al., 2017; Hanemaaijer et al., 2020; Martinello et al., 2015; Yu et al., 2010). To determine the relative contributions of different calcium channel types during AP-evoked calcium influx in mouse prefrontal pyramidal cells, we made whole-cell current-clamp recordings from layer 5 pyramidal neurons in slices prepared from mice aged 20-30 days old. Neurons were filled via whole cell dialysis with an internal solution containing Alexa 594 for morphological identification and the low-affinity calcium indicator Fluo-5F. Three action potentials (APs) were evoked by somatic current injection (1–1.5 nA, 5 ms duration, 20 ms inter-AP interval), and resultant AIS calcium influx was imaged in linescan mode ∼30 µm from the axon hillock (**Fig. 1A**). AP-evoked calcium transients were stable over repeated linescan sets performed at time intervals used for subsequent pharmacological studies (**Fig. 1B, 1D-E**, median normalized peak ΔG/G_sat_ = 92.6% of baseline, IQR = 88.4–104.7%, n = 32). This calcium influx was largely blocked by a cocktail of Ca_V_ antagonists that included blockers of Ca_V_2.1, Ca_V_2.2, Ca_V_2.3, and Ca_V_3 channels (1 µM ω-conotoxin MVIIC, 1 µm ω-conotoxin GVIA, 0.2 µM agatoxin TK, 0.5 µM SNX-482, 2 µM TTA-P2) (**Fig. 1B, 1D**, median normalized peak ΔG/G_sat_ = 34.6%, IQR = 22.9– 48.4%, n = 7, p = 0.002).

Specific channel antagonists were then applied one-by-one to examine contributions from individual Ca_V_ classes. Consistent with previous reports across a range of cell types (Bender and Trussell, 2009; Clarkson et al., 2017; Fukaya et al., 2018; Hanemaaijer et al., 2020; Martinello et al., 2015), Ca_V_3 channels were a major source of AIS calcium influx, as the specific antagonist TTA-P2 reduced calcium influx to 76.6% of baseline (**Fig. 1C, 1E**, IQR = 75.0–84.9%, n = 11, p = 0.002, Mann Whitney U-Test). Additional contributions were made from Ca_V_2 channels, with the Ca_V_2.3-preferring antagonist SNX-482 reducing AIS calcium influx to 77.6% of baseline (**Fig. 1C, 1E**, 500 nM; IQR = 72.4–88.4%, n = 9, p = 0.006). Application of the Ca_V_2.1 channel antagonist ω-agatoxin-TK (200 nM) resulted in variable blockade, with AIS calcium unaffected in some cells and reduced ∼30% in others (**Fig. 1E**, median: 74.1% of baseline, IQR = 68.6– 91.9%, n = 6, p = 0.079). The Ca_V_2.2 antagonist ω-conotoxin GVIA (1 µM) had little to no effect on AIS calcium (**Fig. 1E**, median = 88.9%, IQR = 85.6–91.3%, n = 6, p = 0.133), but the dual Ca_V_2.1/2.2 antagonist ω-conotoxin MVIIC (1 µM) appeared to have an additive effect, blocking ∼40% of total calcium influx (**Fig. 1C, 1E**, median normalized peak ΔG/G_sat_ = 61.5%, IQR = 58.3–73.0%, n = 9, p = 6.19 × 10^−6^). The presence of each of these Ca_V_2.1/2.2 antagonists at the slice was confirmed by monitoring progressive blockade of evoked EPSPs elicited by a glass bipolar stimulating electrode placed 200 µm lateral to the soma in layer 5 (**Table 1**). Lastly, we applied the Ca_V_1 antagonist nifedipine (10 µM), which had no effect on AIS calcium influx (**Fig. 1E**, median = 99.5%, IQR = 91.9–102.3%, n = 7, p = 0.314). We observed no change in action potential peak, threshold, or half-width throughout the recordings (**Table 1**). Together, these data indicate that AP-evoked calcium influx in mouse prefrontal pyramidal cells is supported by a mix of Ca_V_2 and Ca_V_3 channels.

**Table 1:**
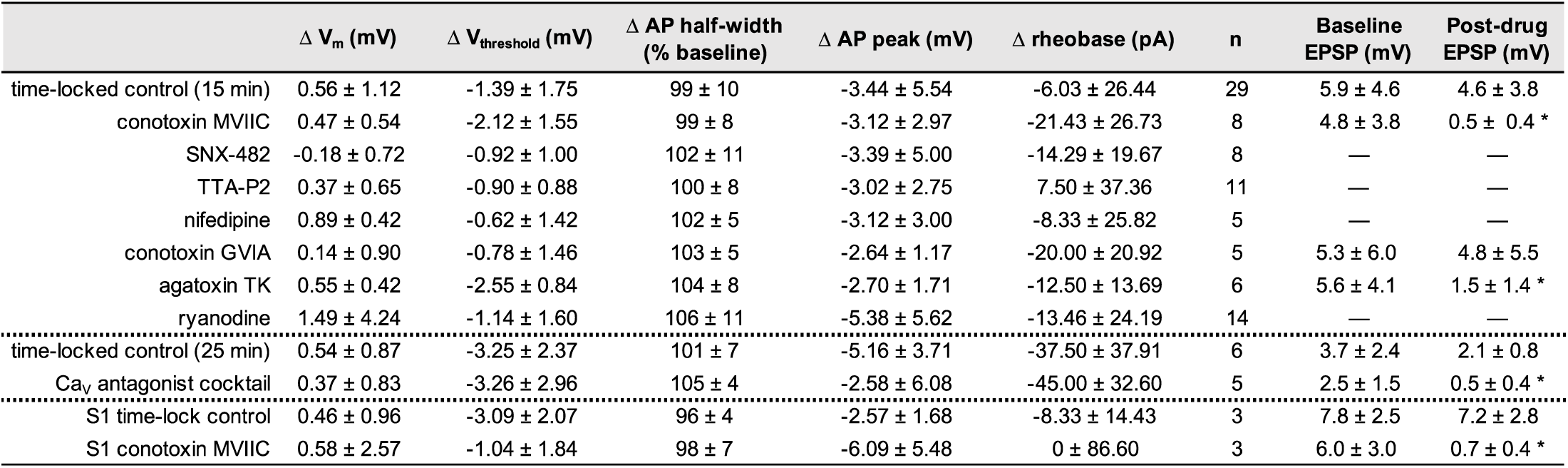
Changes in action potential waveform properties across the course of recording. * denotes a p-value < 0.05. One-way ANOVAs or two-tailed t-tests were performed for each waveform property, as appropriate. Paired t-tests were performed for EPSP amplitudes.

| | $\Delta V_m$ (mV) | $\Delta V_{\text{threshold}}$ (mV) | $\Delta$ AP half-width (% baseline) | $\Delta$ AP peak (mV) | $\Delta$ rheobase (pA) | n | Baseline EPSP (mV) | Post-drug EPSP (mV) |
| --- | --- | --- | --- | --- | --- | --- | --- | --- |
| time-locked control (15 min) | 0.56 ± 1.12 | -1.39 ± 1.75 | 99 ± 10 | -3.44 ± 5.54 | -6.03 ± 26.44 | 29 | 5.9 ± 4.6 | 4.6 ± 3.8 |
| conotoxin MVIIC | 0.47 ± 0.54 | -2.12 ± 1.55 | 99 ± 8 | -3.12 ± 2.97 | -21.43 ± 26.73 | 8 | 4.8 ± 3.8 | 0.5 ± 0.4 * |
| SNX-482 | -0.18 ± 0.72 | -0.92 ± 1.00 | 102 ± 11 | -3.39 ± 5.00 | -14.29 ± 19.67 | 8 | — | — |
| TTA-P2 | 0.37 ± 0.65 | -0.90 ± 0.88 | 100 ± 8 | -3.02 ± 2.75 | 7.50 ± 37.36 | 11 | — | — |
| nifedipine | 0.89 ± 0.42 | -0.62 ± 1.42 | 102 ± 5 | -3.12 ± 3.00 | -8.33 ± 25.82 | 5 | — | — |
| conotoxin GVIA | 0.14 ± 0.90 | -0.78 ± 1.46 | 103 ± 5 | -2.64 ± 1.17 | -20.00 ± 20.92 | 5 | 5.3 ± 6.0 | 4.8 ± 5.5 |
| agatoxin TK | 0.55 ± 0.42 | -2.55 ± 0.84 | 104 ± 8 | -2.70 ± 1.71 | -12.50 ± 13.69 | 6 | 5.6 ± 4.1 | 1.5 ± 1.4 * |
| ryanodine | 1.49 ± 4.24 | -1.14 ± 1.60 | 106 ± 11 | -5.38 ± 5.62 | -13.46 ± 24.19 | 14 | — | — |
| time-locked control (25 min) | 0.54 ± 0.87 | -3.25 ± 2.37 | 101 ± 7 | -5.16 ± 3.71 | -37.50 ± 37.91 | 6 | 3.7 ± 2.4 | 2.1 ± 0.8 |
| Ca<sub>v</sub> antagonist cocktail | 0.37 ± 0.83 | -3.26 ± 2.96 | 105 ± 4 | -2.58 ± 6.08 | -45.00 ± 32.60 | 5 | 2.5 ± 1.5 | 0.5 ± 0.4 * |
| S1 time-lock control | 0.46 ± 0.96 | -3.09 ± 2.07 | 96 ± 4 | -2.57 ± 1.68 | -8.33 ± 14.43 | 3 | 7.8 ± 2.5 | 7.2 ± 2.8 |
| S1 conotoxin MVIIC | 0.58 ± 2.57 | -1.04 ± 1.84 | 98 ± 7 | -6.09 ± 5.48 | 0 ± 86.60 | 3 | 6.0 ± 3.0 | 0.7 ± 0.4 * |

### Ca_V_3 channels couple to ryanodine-dependent stores at the AIS

Calcium-containing cisternal organelles are found in pyramidal cell initial segments throughout the neocortex (Antón-Fernández et al., 2015; Benedeczky et al., 1994; Sánchez-Ponce et al., 2012; Schlüter et al., 2017), but their role as a potential source of calcium during APs is not well understood. These cisternal organelles localize to discrete sites within the AIS of pyramidal cells (King et al., 2014; Schneider-Mizell et al., 2020) and express ryanodine receptors (RyR) which gate calcium-induced calcium release (Chamberlain et al., 1984; Endo et al., 1970; Van Petegem, 2012). Thus, they may boost AP-evoked calcium transients if they are coupled to Ca_V_s in the AIS. To determine whether calcium release from cisternal organelles is recruited at the AIS during AP generation, we began by comparing AP-evoked calcium influx at the AIS before and after ryanodine application, which at high concentrations (>10 µM) blocks calcium-induced calcium release by preventing the opening of ryanodine receptors (Thomas and Williams, 2012). In contrast to somatosensory cortex layer 5b pyramidal neurons, where calcium stores account for ∼50% of AP-evoked calcium transients (Hanemaaijer et al., 2020), ryanodine (20 µM) had a more modest effect in prefrontal cortex, reducing AP-evoked calcium transients to 85.4% of baseline (**Fig. 2A-B**, IQR = 79.2–89.4%, n = 17, p = 0.008). These ryanodine-dependent stores appear to be the sole source of intracellular calcium in the AIS, as subsequent application of the SERCA-ATPase inhibitor cyclopiazonic acid (CPA, 20 µM), which completely depletes calcium stores, did not lead to further decrements in AP-evoked calcium transients. (**Fig. 2A, 2C**, ryanodine: 86.4% of baseline, IQR = 85.0– 89.1%, ryanodine + CPA (30-min application): 77.0% of baseline, IQR = 75.8–82.7%, n = 7, p = 0.108, Wilcoxon Signed-Rank Test). This suggests that ryanodine receptors govern the majority of store-related calcium release during AP activity in the AIS.

Ryanodine receptors can be coupled tightly to Ca_V_s, either through direct physical coupling or through indirect nanodomain proximity (Irie and Trussell, 2017; Johenning et al., 2015). In the AIS, ryanodine-dependent signaling is also important for D3 dopamine receptor-dependent regulation of Ca_V_3s (Yang et al., 2016). To test if ryanodine-dependent stores were preferentially coupled to particular Ca_V_ classes present at the AIS, we performed sequential application of a selective Ca_V_ antagonist followed by ryanodine (20 µM). With this approach, occlusion of any ryanodine-mediated reductions in AIS calcium would suggest that the blocked Ca_V_ was the source of calcium that induced subsequent RyR-dependent store release. Interestingly, we found that block of Ca_V_3 with TTA-P2 produced the clearest occlusion (**Fig. 3A-B**, TTA alone: median = 77.6%, IQR = 74.7–85.5%, TTA plus ryanodine = 72.5%, IQR = 71.2–78.9%, n = 6, p = 0.53, Wilcoxon Signed-Rank Test). Conversely, application of ryanodine after pre-application of ω-conotoxin MVIIC resulted in a significant reduction in AIS calcium (**Fig. 3A-B**, conotoxin alone: median = 65.4%, IQR = 59.7– 74.3%, conotoxin plus ryanodine: median = 56.1%, IQR = 47.4–62.8%, n = 7, p = 0.03, Wilcoxon Signed-Rank Test). An intermediate phenotype was observed with Ca_V_2.3 block by SNX-482; decrements in calcium influx after ryanodine were observed in some cells, but the overall change was not significant (**Fig. 3A-B**, SNX alone: median = 73.8%, IQR = 70.3– 89.8%, SNX plus ryanodine: median = 69.8%, IQR = 52.9–87.8%, n = 7, p = 0.20, Wilcoxon Signed-Rank Test). Overall, these data indicate that, of all Ca_V_ classes found in the AIS, Ca_V_3s are most likely to be in close proximity to cisternal organelles to evoke release of calcium stores.

### Functional distribution of Ca_V_3 and Ca_V_2.1/2.2 in the AIS

Ryanodine receptors are localized to discrete, ankyrin-G deficient regions of the AIS (King et al., 2014). Given the tight association between Ca_V_3 channels and RyR-dependent release, we hypothesized that Ca_V_3 channels may exhibit similar clustering at the functional level, which could be observed using approaches for resolving nanodomain “hotspots” of calcium. Such approaches have been utilized to examine discrete sites of calcium incursion at presynaptic terminals using confocal microscopy entry (DiGregorio et al., 1999; Nakamura et al., 2015), but, to our knowledge, have not been applied at the AIS with two-photon imaging.

To test whether there are sites within the AIS that are hotspots for calcium entry, the excitation laser was parked at one of 5 sites along the wall of the AIS membrane, each 500 nm apart, and APs were evoked while imaging calcium influx at 20 kHz. Calcium influx was reported with a recording solution containing the low-affinity calcium indicator Oregon Green BAPTA-5N (600 µM) supplemented with the slow calcium chelator EGTA (0.1 µM) to restrict imaged signals to sites experiencing rapid, high concentrations of calcium incursion (DiGregorio et al., 1999). Data were quantified by comparing isochronal influx amplitude at the peak within the set of 5 points to the point (or average of points) 1 µm away on either flank. Using this approach, we identified a range of responses, from small differences across all five sites, to areas where certain locations had calcium incursions that were elevated relative to neighboring sites.

In initial experiments that averaged over 50 trials, we found that the majority of sites (46 of 59) fell within a normal distribution (**Fig. 4C**), with no appreciable difference in peak calcium influx across all 5 sites. But in the remainder (13 of 59), calcium influx appeared to be more elevated and have sharper kinetics, consistent with a hotspot for calcium entry. Indeed, calcium entry at these sites was ∼2x larger than in non-hotspot regions (**Fig. 4D**), while flanks 1 µm from the peak both more proximal or more distal to the axon hillock were of comparable amplitude to non-hotspot regions (**Fig. 4E**).

These hotspots may represent sites of concentrated calcium influx through Ca_V_s or reflect coupling to intracellular calcium stores. To test this, imaging was repeated (averages of 20 scans) in the presence of Ca_V_3 antagonists, Ca_V_2.1/2.2 antagonists, or with stores depleted with CPA. Both TTA-P2 and conotoxin-MVIIC reduced overall calcium influx by 33.6% and 17.1%, respectively, whereas CPA had no effect on peak amplitude when compared to control sets acquired with identical approaches (**Fig. 4F**). This is consistent with the hypothesis that such imaging approaches using high concentrations of calcium indicators supplemented with calcium buffers may uncouple Ca_V_-mediated influx from intracellular stores (Collier et al., 2000).

To determine whether Ca_V_3 or Ca_V_2.1/2.2 channels preferentially contribute to hotspot regions, amplitudes at the peak were compared to isochronal amplitudes 1 µm lateral for all data. To determine hotspot frequency after pharmacological manipulation, hotspots were defined as any set with a 1.5x difference between the peak and 1 µm lateral amplitudes. While CPA had no effect on hotspot frequency, application of Ca_V_ antagonists changed hotspot frequency dramatically. TTA-P2 eliminated hotspots almost entirely, whereas conotoxin-MVIIC increased the fraction of observed hotspots (**Fig. 4G**). Taken together, these data indicate that Ca_V_3 channels are uniquely clustered in the AIS, producing nanodomains of elevated calcium entry that then couple to RyR-dependent stores.

### Temporally distinct AP-evoked sodium and calcium dynamics in the AIS

Imaging data above suggests that AIS calcium entry occurs during AP repolarization. While this is consistent with Ca_V_ activity during APs in a range of imaging and electrophysiological studies at various sites within the axon (Bischofberger et al., 2002; Díaz-Rojas et al., 2015; DiGregorio et al., 1999; Nakamura et al., 2015; Rowan et al., 2014), it has recently been proposed that AIS calcium influx during APs is mediated by Na_V_s, not Ca_V_s (Hanemaaijer et al., 2020). If this is the case, then calcium and sodium influx should occur simultaneously. To test this, we started by comparing AP-evoked sodium and calcium transients using 5.3 kHz linescans that transected the AIS 30 µm from the hillock. Linescans were collected at room temperature (22 °C) to best separate the rising and falling phase of the AP. The low-affinity indicator Fluo-4FF was used for calcium imaging and the most commonly utilized sodium indicator, SBFI, was used for sodium imaging. SBFI reports changes in sodium concentration with a shift in emission spectra, which, with 2-photon excitation sources, is best visualized as a reduction in fluorescence (Bender et al., 2010; Rose et al., 1999). Sodium and calcium transients were fitted with sigmoid functions and event onset was defined as the time at which the sigmoid fit first exceeded the amplitude of baseline root-mean-squared noise (RMS). Similar to previous reports (Hanemaaijer et al., 2020), we found that the rising slope of sodium and calcium transients were comparable (**Fig. 5B**, Na median = 0.07% ΔF/F per s, IQR = 0.06–0.08% ΔF/F per s, n = 11, Ca median = 0.07% ΔF/F per s, IQR = 0.04–0.08% ΔF/F per s, n = 9, p = 0.6, Mann-Whitney); however, sodium influx typically occurred during the rising phase of the AP whereas calcium influx occurred during the falling phase (relative to AP peak, Na median = -0.343 ms, IQR = -0.4045–0.063 ms, n = 11 cells, Ca median = 0.901 s, IQR = 0.622–1.284, n = 9 cells, p = 0.0007, Mann-Whitney). The mean difference in transient onset was 1.2 ms, comparable to the duration of the rising phase of the AP in these recording conditions (median = 0.85 ms, IQR = 0.76–0.94 ms, n = 20 cells).

SBFI typically reported ion influx with a higher signal-to-noise ratio than Fluo-4FF (peak amplitude/baseline RMS). This alone may account for the earlier event onset for SBFI-based signals. To test whether this was the case, we analyzed the subset of data in which signal-to-noise was comparable between sodium and calcium imaging scans (**Fig. 5B**). In these cases, sodium influx still preceded calcium influx. Thus, these data suggest that sodium and calcium influx occur through distinct mechanisms that can be separated temporally.

Previous work has suggested that the timing of calcium influx during an AP may shift to earlier parts of the AP at high temperature, in part due to differences in gating kinetics between Na_V_s and Ca_V_s (Sabatini and Regehr, 1996). This would be best assayed with the temporal fidelity of pointscan imaging. Unfortunately, we found the high basal fluorescence of SBFI resulted in significant photo-toxicity when the laser was parked at single sites. Therefore, we made use of a relatively new sodium-sensitive dye, ING-2, which reports increases in sodium concentration with an increase in fluorescence intensity without a change in emission spectra (Filipis and Canepari, 2020). Sodium influx was imaged in sets of 5 sites each 0.5 µm apart, as done for calcium pointscan imaging. But in contrast to calcium imaging data, sodium influx did not appear to occur with regions that could be defined as hotspots. Rather, data reporting the relative amplitudes of the peak sodium transient relative to a neighbor 1 µm away could all be fit within a normal distribution (**Fig. 6B-C**, Shapiro-Wilk test for normality, p = 0.36), consistent with relatively constant Na_V_ density throughout the AIS (Leterrier, 2018).

Similar to data obtained at 22 °C, sodium influx imaged 25-35 µm from the axon hillock again preceded the peak of the AP at 32-34°C (median = -0.65 ms, IQR = -0.9625– -0.5ms, n = 56 sites, 23 cells). Moreover, these events tended to precede AP onset as measured in the soma (median = -0.05 ms, IQR = -0.3625– 0.1 ms). This may be due in part to the conduction delay between the AIS site of AP initiation and the soma (Kole et al., 2007; Rowan et al., 2014), and in part to subthreshold sodium influx before AP onset (Filipis and Canepari, 2020).

Comparisons with onset kinetics of calcium transients imaged with pointscan approaches again revealed marked differences between the onset of sodium and calcium entry. Calcium influx was detectable 1.2 ms after sodium influx, typically during the first millisecond of AP repolarization (**Fig. 6D**, median = 0.75, IQR =0.65–1.35 ms after AP threshold, median = 0.35, IQR = 0.15–0.85 ms after AP peak, n = 37 sites, 12 cells). Similar results were obtained in conditions where Ca_V_3 or Ca_V_2.1/2.2 channels were blocked (TTA-P2: median = 1.1 ms, IQR = 0.8–1.6 ms after AP threshold, median = 0.6 ms, IQR = 0.3–1.1 ms after AP peak, n = 33 point sets, 7 cells; ω-conotoxin MVIIC: median = 1.25 ms, IQR = 0.9625–1.575 ms after AP threshold, median = 0.7 ms, IQR = 0.375–1.1 ms after AP peak, n = 16 sites, 5 cells), and when intracellular stores were depleted with CPA (median = 0.7 ms, IQR = 0.4–0.9375 ms after AP threshold, median = 0.15 ms, IQR = -0.0375–0.4875 ms after AP peak, n = 20 point sets, 5 cells).

Overall, these data are most consistent with the hypothesis that Ca_V_s are the sole source of calcium influx from the extracellular space in the AIS. This contrasts with work in rat somatosensory cortex (S1), where AP-evoked calcium transients were partially blocked by the Ca_V_3 antagonist TTA-P2 (1 µM at equilibrium in the extracellular solution), but, notably, not affected by the Ca_V_2 peptide antagonist ω-conotoxin MVIIC (2 µM, applied via pressure ejection local to the AIS) (Hanemaaijer et al., 2020). Therefore, we tested whether mouse S1 pyramidal cells differ from mouse prefrontal pyramidal cells in the expression of Ca_V_2.1/2.2 channels in the AIS by applying ω-conotoxin MVIIC (1 µM at equilibrium in the extracellular solution) to L5 pyramidal neurons in the somatosensory cortex, again imaging calcium influx resulting from 3 APs. Similar to mouse prefrontal cortex, block of Ca_V_2.1/2.2 channels reduced peak calcium influx by over 35% (**Fig. 7A-B**, median = 63.8%, IQR = 42.5–69.9%, n = 5; time-locked controls: median = 90.9%, IQR = 79.6–95.5%, n = 5; p = 0.012, Mann Whitney U-Test). Thus, these data indicate that pyramidal cells in multiple neocortical regions express a mix of Ca_V_2 and Ca_V_3 channels in their initial segments, at least in the mouse brain.

**Figure 7:**
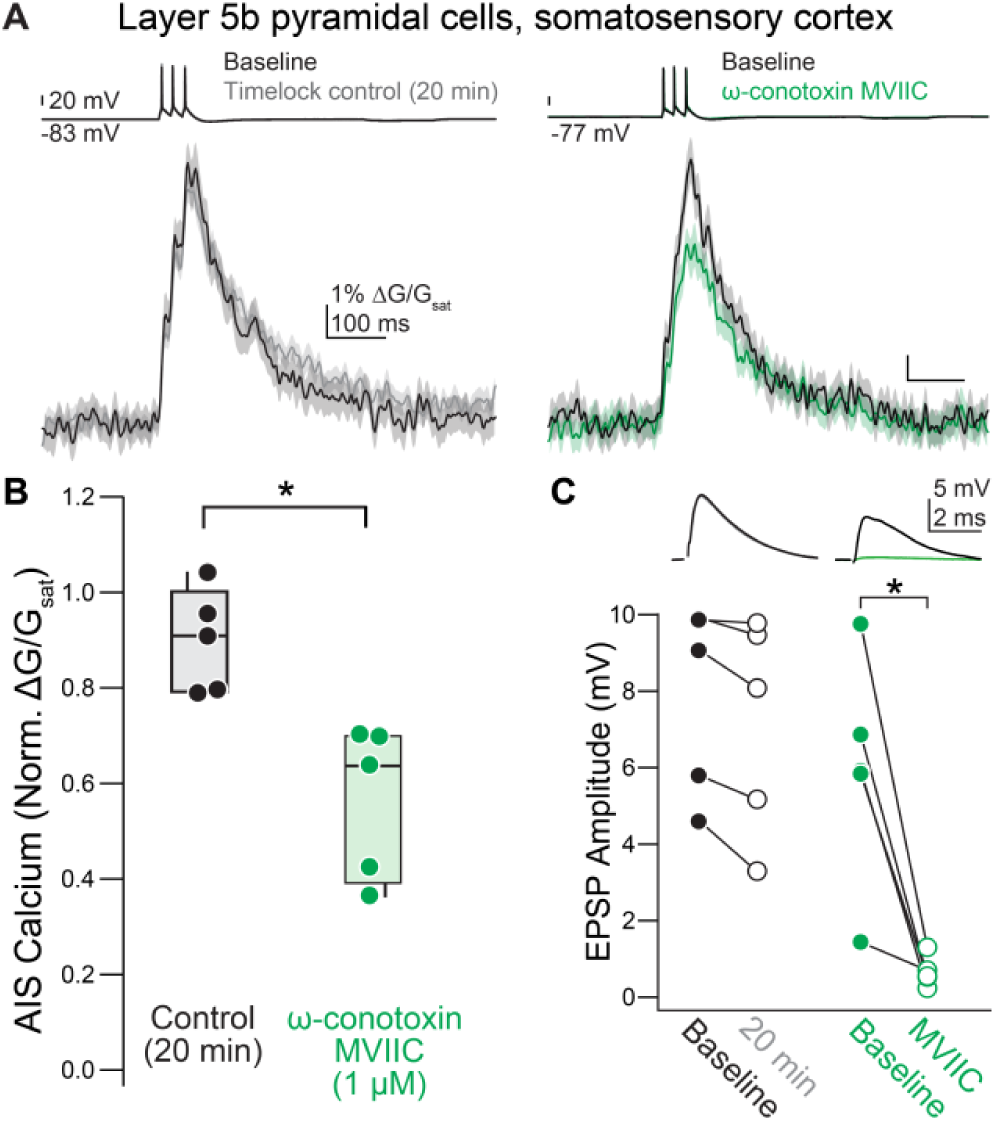
Ca_V_2.1/2.2 contribute to AIS calcium in the somatosensory cortex. A. Representative effect of ω-conotoxin MVIIC application on AIS calcium in L5b pyramidal cells in the somatosensory cortex. Left: example time-locked control cell. Black, baseline; gray, post. Right: example of the effect of ω-conotoxin MVIIC. Black, baseline; green, ω-conotoxin MVIIC. Linescan data are plotted as mean ± standard error. B. Summary of the effects of ω-conotoxin MVIIC on AIS calcium in somatosensory L5b pyramidal neurons. Black, time-locked control cells; green, ω-conotoxin MVIIC. C. Decreases in EPSP amplitude confirm the presence of ω-conotoxin MVIIC at the slice. Top: representative examples ofPSPs in ω-conotoxin MVIIC (right) or in time-locked control cells (left). Bottom: Summary of the effects of ω-conotoxin MVIIC on EPSP amplitude in somatosensory cortex. Black, baseline; gray, time-locked control; green, ω-conotoxin MVIIC.

## DISCUSSION

Here, we show that calcium channels are functionally distributed in distinct domains within mouse prefrontal pyramidal cell initial segments. Low voltage-activated Ca_V_3-mediated calcium influx occurs in spatially restricted “hotspots” whereas high voltage-activated Ca_V_2.1 and Ca_V_2.2 channels provide a more diffuse source of calcium. Ca_V_3-mediated hotspots preferentially evoked additional calcium release from ryanodine-dependent intracellular stores, suggesting that hotspots of Ca_V_3-mediated influx localize to regions enriched with RyRs that are also sites for GABAergic input and K_V_2.1 clustering (King et al., 2014). This suggests that different Ca_V_ classes are functionally localized to discrete regions int the AIS.

### Activity-dependent calcium sources in the AIS

Though AP-evoked calcium influx at the AIS is well-established (Callewaert et al., 1996; Schiller et al., 1995), the sources of this calcium influx have only been investigated relatively recently. These sources appear to be remarkably heterogeneous across neuronal classes and species. In mouse auditory brainstem cartwheel cells, Ca_V_3.2 and SNX-sensitive Ca_V_2.3 account for ∼90% of AP-evoked calcium influx, with no contributions from Ca_V_2.1 or Ca_V_2.2 (Bender and Trussell, 2009). By contrast, the first study of pyramidal cells in ferret neocortex found that calcium influx was mediated by Ca_V_2.1 and Ca_V_2.2, but not Ca_V_3 (Yu et al., 2010). Here, we find that prefrontal pyramidal cells in mouse prefrontal cortex exhibit Ca_V_3-mediated influx, consistent with previous reports in rodent neocortex (Clarkson et al., 2017; Hanemaaijer et al., 2020) and other brain regions (Gründemann and Clark, 2015; Jin et al., 2019; Martinello et al., 2015). Ca_V_2.1/2.2 and Ca_V_2.3 were also found to contribute to calcium influx, highlighting the relative complexity of calcium signaling in prefrontal pyramidal cell initial segments.

In addition to Ca_V_-mediated calcium influx, we found that a small fraction of AP-evoked calcium was released from ryanodine-dependent intracellular stores in the AIS. Cisternal organelles at the AIS were proposed to be involved in calcium sequestration due to their expression of a calcium pump (Ca^2+^-ATPase) in pyramidal neurons of the hippocampus (Benedeczky et al, 1994). Cisternal organelles were originally identified in the initial segments of cortical principal neurons in sensory cortical regions (Benedeczky et al., 1994; Peters et al., 1968). In these regions, a subpopulation of layer 5 pyramidal neurons contain a giant saccular organelle that extends through the entire AIS and accounts for a major fraction of AP-evoked calcium signals (Antón-Fernández et al., 2015; Hanemaaijer et al., 2020; Sánchez-Ponce et al., 2012). Subsequent work has implicated both RyR-dependent and inositol 1,4,5-triphosphate (IP_3_) receptor-dependent AIS-localized stores in a range of processes, including calcium influx during APs, modulation of AIS-associated proteins, and experience-dependent structural plasticity of the AIS compartment (Gomez et al., 2020; Irie and Trussell, 2017; Schlüter et al., 2017; Yang et al., 2016). These different effects may reflect diverse structures and functions in AIS calcium stores across cell types. Conversely, different modes of calcium release from intracellular stores may be recruited by different stimuli.

A recent study in rat somatosensory cortex found that roughly 75% of AP-evoked calcium signaling in the AIS was mediated by a mix of Ca_V_3-mediated influx and release from intracellular stores (Hanemaaijer et al., 2020). Of note, peptide antagonists of Ca_V_2.x channels, puffed for 3s onto the AIS, did not affect AIS calcium influx, despite almost completely blocking AP-evoked calcium signals in axonal boutons. It was therefore proposed that the residual influx was through Na_V_s rather than Ca_V_s, based in part on the observations that influx was sensitive to Na_V_ antagonists and that the rising kinetics of sodium and calcium transients were similar (Hanemaaijer et al., 2020). By contrast, we observed a marked block of AP-evoked influx in prefrontal and somatosensory pyramidal cells from Ca_V_2 antagonists when allowed to equilibrate in the extracellular solution (**Fig. 1, Fig 7**). Furthermore, we found that the kinetics of sodium and calcium influx were indeed identical, but that calcium influx lagged sodium influx in ways that were consistent with sodium and calcium influx occurring on the rising and falling phases of the AP, respectively. These results are consistent with studies using high-speed optical imaging, where sodium influx occurs during the rising phase of the AP (Filipis and Canepari, 2020), whereas calcium influx occurs during the falling phase of the AP in the AIS or AIS-like regions of AP initiation (Hanemaaijer et al., 2020; Pressler and Strowbridge, 2019). Furthermore, we observed consistent results at both room temperature and physiological temperatures with two different sodium-sensitive indicators and two different calcium-sensitive indicators, suggesting that calcium influx occurs during Ca_V_-mediated tail currents on the falling phase of the AP in the axon, regardless of temperature (Kawaguchi and Sakaba, 2015; Pressler and Strowbridge, 2019; Rowan et al., 2014; Sabatini and Regehr, 1996). Nevertheless, calcium influx, as assayed with synthetic indicators, could not be blocked completely with Ca_V_ antagonists. This may be due to several issues, including incomplete block of Ca_V_2.3, or R-type calcium channels, so named for their resistance to antagonist block. Indeed, careful pharmacological studies across cell classes have shown that Ca_V_2.3 channels in pyramidal cells are particularly resistant to block by SNX-482 (Sochivko et al., 2002). Furthermore, block of Ca_V_s by peptide toxins can have relatively slow kinetics (McDonough et al., 1996), and while we made every effort to allow for equilibration, with application times exceeding 20 min, this may not have been sufficient for complete block. Regardless, the kinetics of AIS calcium transients, observed here and in other reports (Hanemaaijer et al., 2020; Pressler and Strowbridge, 2019), are most consistent with influx predominantly through Ca_V_s.

### Functional compartmentalization of calcium influx within the AIS

In mature neocortical pyramidal cells, Na_V_1.6 channels cluster in the regions of the AIS more distal to the soma, whereas Na_V_1.2 channels cluster in the region more proximal to the soma (Hu et al., 2009). This subcompartmental distribution affects the integrative properties of the AIS in health and disease (Hu et al., 2009; Spratt et al., 2019), and raised the question of whether similar functional specializations are found in Ca_V_s localized to the AIS. To test this, we adapted spot imaging techniques used previously to observe calcium microdomains with single-photon sources for use with 2-photon microscopy (DiGregorio et al., 1999; Nakamura et al., 2015). This approach revealed that calcium influx in the AIS occurs in two domains, with hotspots of calcium interspersed within regions of more consistent calcium influx (**Fig. 4**). These calcium nanodomains are hypothesized to result from channel clustering, as isochronal calcium measurements at increasing distances from the calcium source decreased in amplitude, a consequence of calcium diffusing away from its entry site (DiGregorio et al., 1999). It is plausible that the hotspots observed here represent points that are, by chance, closer to clusters of Ca_V_s; however, the differential pharmacological block of hotspots and non-hotspots with selective Ca_V_ antagonists indicates that these hotspots indeed reflect a differential organization of Ca_V_ channel types at the AIS. In future efforts, it will be important to develop immunostaining methods sensitive enough to visualize these channels relative to other AIS constituents to validate these functional observations.

The biophysics of different Ca_V_ channel types may shape calcium hotspot kinetics and duration as well. Relative to currents measured by step-commands, proportionally more current is carried by low voltage-activated than high voltage-activated channels during an AP waveform (McCobb and Beam, 1991). Low voltage-activated channels, including Ca_V_3, open earlier in the course of the AP, and, due to their slower deactivation kinetics, remain open longer than high voltage-activated channels, resulting in a longer duration of calcium influx through these channels (Lambert et al., 1998; McCobb and Beam, 1991). Hotspot calcium influx observed here is consistent with these biophysical aspects of AP-evoked Ca_V_3-mediated currents.

Pharmacological block of ryanodine-dependent stores indicates that Ca_V_3s preferentially couple to intracellular sources of calcium in the AIS, which are found at ankyrin-G deficient regions of the axonal membrane (King et al., 2014). The components of the cytoskeletal scaffolding machinery that tether Na_V_, K_V_ channels, and GABA_A_ receptors in the AIS have been well-characterized (Leterrier, 2018), but how Ca_V_s are anchored at the AIS remains an open question. One possibility, at least for Ca_V_3s clustered with RyRs, are K_V_2.1 channels. K_V_2.1 channels have been shown to tether Ca_V_1 channels near junctions between the endoplasmic reticulum and plasma membrane (Fox et al., 2015), but whether or not they tether Ca_V_s near the cisternal organelle at the AIS has not been explored. Another candidate is amphiphysin II/Bridging integrator 1 (BIN1), a T-tubule protein involved in localizing Ca_V_1.2 channels in cardiac myocytes (Hong et al., 2010). This protein shows specific localization to neuronal AIS and nodes of Ranvier, but whether this protein interacts with AIS channels has not been explored (Butler et al., 1997). Additionally, the presence of auxiliary subunits on Ca_V_1 and Ca_V_2 channels has been shown to affect localization and membrane expression (Arikkath and Campbell, 2003). As Ca_V_3 channels do not associate with auxiliary subunits (Simms and Zamponi, 2014), Ca_V_3 and Ca_V_2 could acquire differential expression within the AIS through differential association of auxiliary subunits with scaffolding elements.

### Functional implications of calcium channel compartmentalization within the AIS

GABA_A_ receptors cluster in ankyrin-G deficient pockets of the AIS and associate with clustered non-conducting K_V_2.1 channels that stabilize junctions between cisternal organelles and the plasma membrane (Benedeczky et al., 1994; King et al., 2014; Kirmiz et al., 2018; Schneider-Mizell et al., 2020). These clustering domains appear across species and brain regions (King et al., 2014). The coupling of Ca_V_3 channels to ryanodine receptors, as well as the clustering of these channels into hotspots, suggests that Ca_V_3 channels co-localize with GABAergic chandelier synapses in the AIS. Thus, AIS Ca_V_3s may be particularly sensitive to chandelier cell input. In mature neurons, hyperpolarizing inhibition has been shown to relieve Ca_V_3 channels from steady-state inactivation, thereby promoting rebound spike bursts immediately following an inhibitory epoch (Molineux et al., 2006; Ulrich and Huguenard, 1997). Interestingly, chandelier inputs switch from depolarizing to hyperpolarizing the AIS membrane relatively late in development (Lipkin and Bender, 2020; Pan-Vazquez et al., 2020; Rinetti-Vargas et al., 2017), corresponding to the emergence of synchronized higher-order rhythmicity in cortical networks (Uhlhaas and Singer, 2011). Whether this tight coupling between AIS GABAergic inputs and Ca_V_3s contributes to the development of these network phenomena remains to be explored.

In addition to regulation by chandelier inputs, calcium hotspots could enable precise neuromodulatory control over spike properties, perhaps within select temporal windows relative to neuromodulator signals. In striatal medium spiny neurons, a form of credit assignment for synapses that encode information relevant to reward has been demonstrated based on coincident dopaminergic and glutamatergic signaling (Yagishita et al., 2014). In cells that express D3 dopamine receptors, AIS Ca_V_3 function can be modulated in ways that hyperpolarize the voltage dependence of channel inactivation, in turn lowering the number of channels available for activation during subsequent APs (Clarkson et al., 2017; Yang et al., 2016), This process depends on RyR-dependent intracellular stores (Yang et al., 2016). Thus, in D3 receptor-expressing neurons, Ca_V_3 channels may be modulated only when dopamine binding to D3 receptors coincides with neuronal activity that promotes calcium influx through AIS Ca_V_3s. This may result in preferential suppression of Ca_V_3 function in cells that are actively spiking, thereby modulating only the population of neurons that were active during dopaminergic signaling.

## Author contributions

Conceptualization: AML, MMC, PWES, SLM, KJB. Data curation: AML, KJB. Formal Analysis: AML, KJB. Funding acquisition: AML, PWES, MMC, SLM, KJB. Investigation: AML, MMC, PWES, SLM, KJB. Methodology: AML, KJB. Software: AML, PWES, SLM, KJB. Supervision: KJB. Writing — original draft: AML. Writing — review & editing: AML, MMC, PWES, SLM, KJB.

## Acknowledgements

We are grateful to all members of the Bender Lab who provided input and comments on this manuscript. This work was supported by grants to AML (NSF 1650113), PWES (NSERC PGS-D Scholarship), MMC (NSF 1144247), and KJB (NIH AA027023, MH112729).

